# Decoding cellular deformation from pseudo-simultaneously observed Rho GTPase activities

**DOI:** 10.1101/2021.08.21.457182

**Authors:** Katsuyuki Kunida, Nobuhiro Takagi, Kazushi Ikeda, Takeshi Nakamura, Yuichi Sakumura

**Affiliations:** Graduate School of Science and Technology, Nara Institute of Science and Technology, Ikoma, Nara, 8916-5, Japan; Graduate School of Information Science and Technology, Aichi Prefectural University, Nagakute, Aichi, 480-1342, Japan; Research Institute for Biomedical Sciences, Tokyo University of Science, Noda, Chiba, 278-8510, Japan; Data Science Center, Nara Institute of Science and Technology, Ikoma, Nara, 8916-5, Japan; Fujita Health University School of Medicine, Toyoake, Aichi, 470-1192, Japan

## Abstract

The inability to simultaneously observe all of the important Rho GTPases (Cdc42, Rac1, and RhoA) has prevented us from obtaining evidence of their coordinated regulation during cell deformation. Here, we propose Motion-Triggered Average (MTA), an algorithm that converts individually observed GTPases into pseudo-simultaneous observations. Using the time series obtained by MTA and mathematical model, we succeeded for the first time in decoding the cell edge velocity from the three GTPase activities to provide clear numerical evidence for coordinated cell edge regulation by the three GTPases. We found that the characteristics of the obtained activities were consistent with those of previous studies, and that GTPase activities and their derivatives were involved in edge regulation. Our approach provides an effective strategy for using single-molecule observations to elucidate problems hampered by the lack of simultaneous observations.

## Introduction

Cell migration consists of upstream biochemical reactions, intermediate processes of cytoskeletal changes, and downstream output processes leading to cellular deformation. Among the essential Rho GTPases, which are the main regulators of cytoskeletal reorganization, Cdc42 and Rac1 promote cytoskeletal formation, whereas RhoA is involved in myosin II–mediated cytoskeletal retraction as well as cytoskeletal formation (Etienne-Manneville and Hall, 2002; Hall, 1998; Jaffe and Hall, 2005; Pertz, 2010). This suggests that Rho GTPases regulate the balance between cytoskeletal expansion and retraction at the molecular scale; this idea is phenomenologically consistent with the morphological-scale effects of GTPase activity on cellular events. Recent studies have clarified the upstream signaling of Rho GTPases, including their interactions with guanine nucleotide exchange factors (GEFs) and GTPase-activating proteins (GAPs) (Müller et al., 2020). Cells use these molecules to coordinate cytoskeletal reorganization in a dynamically changing environment. Keeping pace with these investigations, recent bioengineering efforts have yielded biosensors that enable simultaneous live observation of multiple GTPase activities and changes in cell morphology (Machacek et al., 2009; Shcherbakova et al., 2018). This technique provides spatiotemporal data on activities and morphological changes, which could contribute to understanding the coordinated regulation of morphological changes by GTPases (Welch et al., 2011). However, the finite number of optical wavelengths that can be separated also limits the number of GTPases that can be observed simultaneously.

Compared to the huge advances in biosensor development, there has been less progress on technology for data analysis of molecular activity and changes in cell morphology. Hence, there is still no evidence that the three Rho GTPases coordinately regulate cell deformation, i.e., that two of the three Rho GTPases alone are not sufficient, and a fourth factor is not necessary. Many groups have evaluated the relationship between molecular activity and cell edge velocity using temporal cross-correlation (Kunida et al., 2012; Machacek et al., 2009; Tsai et al., 2019). However, there are several drawbacks to this approach. Even though multiple input stimuli drive the dynamical system (**Fig. 1a**), temporal cross-correlation calculates coefficients with a time lag between the time series of only one input and its response time series (**Fig. 1b**). Cross-correlation is an index that averages the similarity of the increase or decrease patterns or the “simultaneity” of the positive and negative peaks between the two time series. Because multiple inputs drive the system’s response, it is impossible to find a causal relationship in the time lags between the response and a single input. This is true largely because the maximum coefficient of temporal cross-correlation is small (almost always less than 0.5). The peak time of the response often precedes that of the input stimulus (**Fig. 1b**), and this is the opposite of the actual causal relationship in time. The fatal flaw in applying temporal cross-correlation to cell migration data is that the correlation coefficients take the time average of the series. Consequently, we lose information on the temporal patterns of molecular activity and edge velocity.

**Figure 1.**
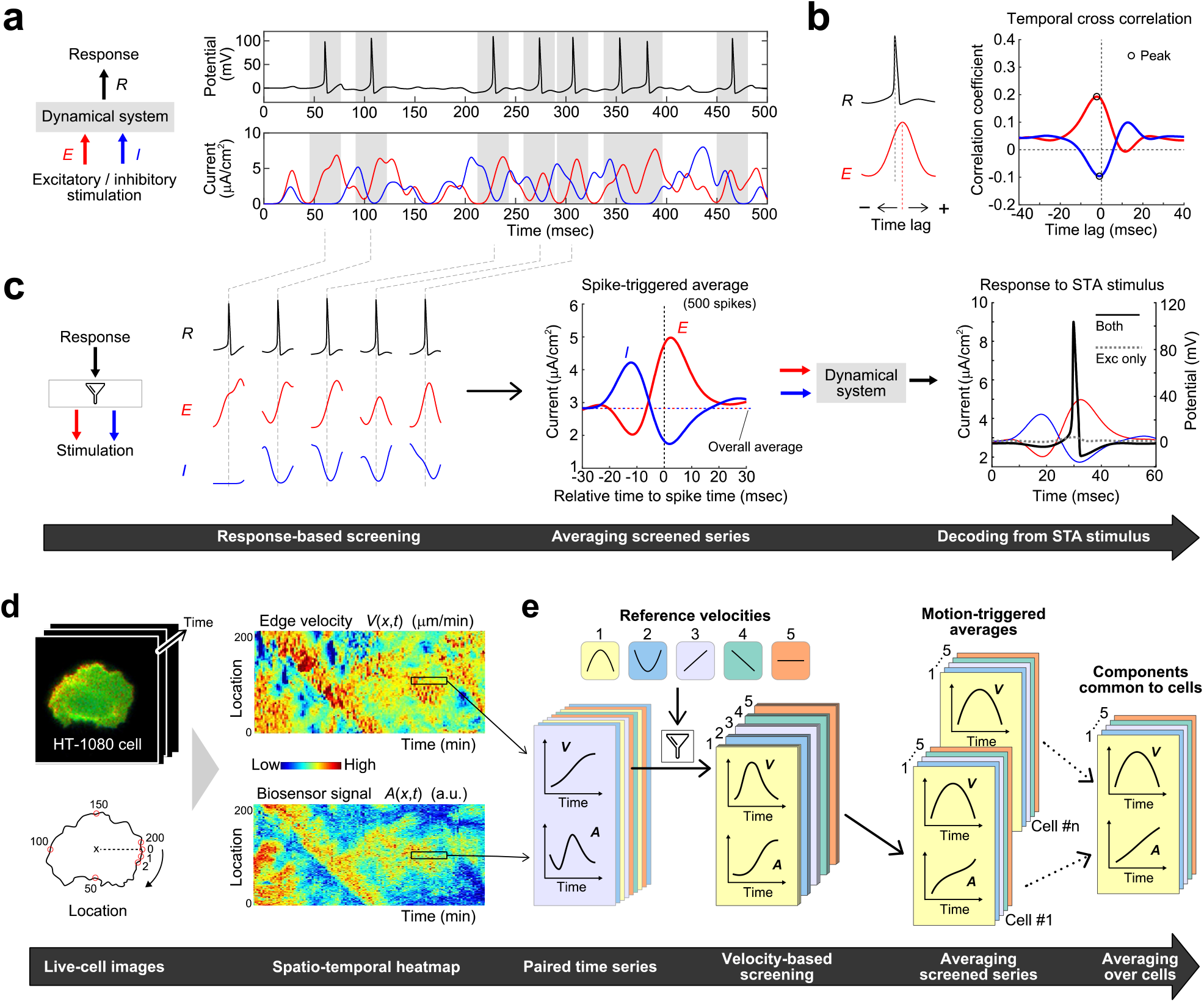
Existing and proposed methods for quantifying relationships between time series of different attributes. **a**. Sample time series obtained from the Hodgkin–Huxley model (Hodgkin and Huxley, 1952). The model receives excitatory (*E*; red curve) and inhibitory (*I*; blue curve) stimuli and responds to them (*R*; black curve). We simulated stimulus currents as Gaussians with random times and amplitudes. **b**. Temporal cross-correlation analysis of the time series in **a**. The stimulation time series was time-shifted into the future (+) and past (−) (left), and temporal cross-correlation calculates correlations calculated between response and excitation (red in the right), and response and inhibition (blue in the right) for each time lag. Black circles indicating maxima and minima lag slightly in the past, meaning that the response reaches its peak before the stimulus. **c**. Spike-triggered average (STA) analysis of the time series in **a** and validation by decoding spike generation. The left shows the partial time series of excitation, inhibition, and response screened around spike peak times (red, blue, and black curves, respectively). The middle panel compares the averages of the screened excitatory and inhibitory time series (solid red and blue lines) with the averages of the overall time series (horizontal dashed line). The right panel confirms that a spike (black line) can be decoded from excitatory (red dashed line) and inhibitory (blue dashed line) stimuli extracted by STA. Only excitatory stimulation generates no spikes (gray dashed line). **d**. Spatio-temporal heatmaps from live-cell images by quantifying edge velocity and molecular activity along the edge of the cell (Kunida et al., 2012). Molecular activity is standardized to a mean of 0 and a standard deviation of 1. **e**. Schematic flowchart of the proposed method, Motion-Triggered Average (MTA), inspired by STA. We prepared pairs of velocity and activity time series in a short time duration (seven frames) without overlapping time or location. We then screened the time-series pairs at a velocity that matched a pre-defined reference velocity pattern. The average of the screened velocities and molecular activities yielded the average activity pattern simultaneously as the edges moved like the reference velocity pattern. We applied MTA to multiple cells with the same reference velocity and averaged the activities over the cells to extract the components common to all cells.

Therefore, in this study, we tried to develop a new analytical method for processing the data obtained by biosensors. The first requirement of the method is preprocessing that converts the individually observed molecular activities of different species into a pseudo-simultaneous dataset. The preprocessing enables us to break through the aforementioned limitations of observation technology. The second requirement is preprocessing to extract GTPase activity components common to multiple cells, observed individually. Spatiotemporal heatmaps of molecular activity and edge velocity (**Fig. 1d**) and clustering of velocity time series (Wang et al., 2018) are helpful for visualizing a single cell’s molecular activity and edge velocity. However, in order to decipher mechanisms and laws that are universal to the target cell types for morphological changes, we must extract the activity components that are common to multiple cells. The final requirement is the validation of the quantitative relationship between activity and velocity extracted by the preprocessing steps. The activity extracted by preprocessing is a necessary condition obtained independently for each GTPase, and we do not know whether the three Rho GTPases acting in coordination are sufficient to regulate the edge velocity. Therefore, we must determine whether we can decode the edge velocity from multiple Rho GTPase activities.

The Spike-Triggered Average (STA) in neuroscience (Bialek et al., 1991; Mainen and Sejnowski, 1995; Marmarelis and Naka, 1972) screens and averages electrical stimulus time series conditioned on the occurrence of neuronal spikes (action potentials) to convert individually observed spikes and multiple stimuli into pseudo-simultaneous observation data (middle panel of **Fig. 1c**). The stimulus time series obtained by STA are at least necessary for spike generation, but we do not know if they contain sufficient information for spike generation. By applying those time series to the original dynamical system, we can verify that they are sufficient to produce spikes and that a single stimulus is not (right panel of **Fig. 1c**). The advantage of STA is that it can integrate data and extract components common to neurons using a spike onset time as a reference signal, even under severe conditions such as random input stimuli, observations of different neurons, and individual observations of each excitatory and inhibitory stimulus. Thus, the STA extracts the property that real neurons, which in this context can be thought of as black boxes, are more sensitive to the amount of change than the magnitude of the input stimulus (Mainen and Sejnowski, 1995).

The essence of STA is the integration of multiple events based on temporal simultaneity. In this study, we propose the Motion-Triggered Average (MTA) algorithm, which screens and averages the molecular activity time series coinciding with the cell edge’s specific pattern velocity time series. Using MTA, we can convert the individually observed molecular activities and edge velocities into a pseudo-simultaneous dataset. MTA also offsets cell-to-cell variation by averaging and extracting common cellular components. We applied MTA with five different reference velocity patterns, including expansion and retraction, to live-cell images of Cdc42, Rac1, and RhoA observed individually in a previous study (Kunida et al., 2012). Thus, we extracted the average activity patterns of each of the three GTPases, coinciding with a velocity pattern similar to each reference velocity, and compared them to those reported in previous studies. Next, we investigated whether these average activity time series could be responsible for the corresponding velocity patterns. There is no quantitative mathematical model of molecular activity–induced edge velocity, such as the neuronal model in **Fig. 1c**. Therefore, we constructed a function of GTPase activities for edge velocity that considered the cell membrane’s elastic properties and performed regression and model selection on the time series obtained by MTA. As expected, we confirmed that the optimal model can decode the correct edge velocity time series from an unknown MTA activity time series. If decoding is possible, the three GTPase activities obtained by MTA will provide numerical evidence for the dynamic coordinated regulation of edge velocity, and the model will represent the decoding strategy.

## Methods and Results

### The Motion-Triggered Average algorithm extracts GTPase activity patterns specific to velocity patterns

MTA is an averaging system that screens time series based on velocity patterns. First, we create spatio-temporal heatmaps of edge velocity and molecular activity from live-cell imaging data (**Fig. 1d**). The activity is standardized to a mean of 0 and a standard deviation of 1. We provide short time series pairs of velocity and activity at the same location and time in the heatmap and screen the paired series with velocity patterns that match the reference velocity patterns prepared in advance. Then, by averaging these screened paired series for velocity and activity, we extract the average series of molecular activity coinciding with the specific velocity pattern of the edge (**Fig. 1e**; **Fig. S1**). If the molecular activity is related to velocity regulation, it will change in time as the velocity changes; otherwise, the MTA activity will be a series of constant values because MTA averages out the equivalent noise at any point in time. Finally, we extract the components common to cells by averaging over multiple cells and offsetting the variation component caused by cell characteristics. By applying the above procedure to each member of Rho GTPases, we can obtain the activities of any individually observed molecules pseudo-simultaneously. This study used live-cell images of three Rho GTPase activities (Cdc42, n = 4; Rac1, n = 4; RhoA, n = 7) measured by FRET imaging (Kunida et al., 2012). We used a total of 15 reference velocity patterns, which consisted of five different patterns (“Expansion,” “Retraction,” “Acceleration,” “Deceleration,” and “Constant”) with seven observation time points and their three levels of intensity (low, middle, and high). These reference velocities were from different time series and classified most of the observed edge velocities (**Fig. S2**).

MTA extracted the velocity-dependent activity time series for each of the three Rho GTPases (**Fig. 2**). To compare the activity levels after the initial time, we displayed the time series with the relative value of zero activity at the initial time (see **Fig. S3** for the original series). We tested the reproducibility, specificity, and generality of these time series. First, to verify the reproducibility of the MTA activity patterns obtained from each of the multiple cells, we examined the cell-to-cell variation in the time series (**Fig. S3**). Despite the small cell number, the mean activities had a slight variation (S.E.), indicating that different cells had reproducible activity levels for the same reference velocity. Each type of Rho GTPase exhibited different activity time series that varied smoothly with significant non-zero values if the velocity was not zero. Therefore, we can regard the activity time series averaged across cells as components common to all cells, without cell-specific variations. In addition, the order of the activity patterns has the same topology as that of the three velocity intensities (in the order blue, yellow, red), suggesting that the activity levels are likely to represent information about velocity.

**Figure 2.**
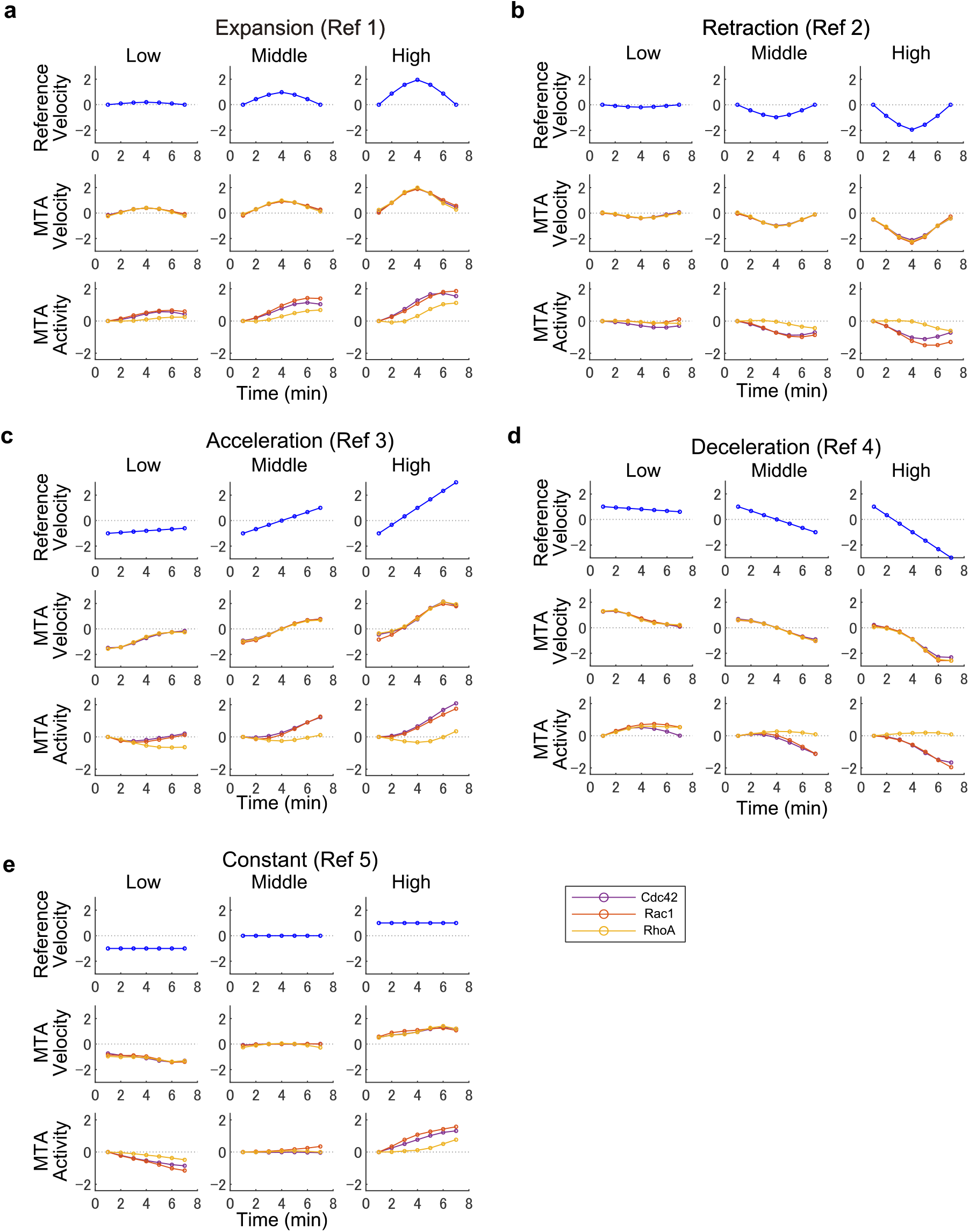
Velocities and GTPase activities referring to five reference velocity patterns using MTA analysis. **a**–**e**. MTA time series for each of the five reference velocity series, (**a**) “Expansion,” (**b**) “Retraction,” (**c**) “Acceleration,” (**d**) “Deceleration,” and (**e**) “Constant.” The MTA velocity series approximates the reference velocity series, and the MTA activity series of Cdc42 (purple), Rac1 (orange), and RhoA (yellow) represent their unique characteristics. We displayed the time series with a relative value of zero activity at the initial time (see **Fig. S3** for actual activity values).

Next, we investigated whether the activity time series were specific to the edge velocity patterns (**Fig. S4**). The MTA time series were constant near zero for activity time series unrelated to the velocity patterns. We performed MTA analysis on a negative control dataset with probes unrelated to velocity regulation (Kurokawa and Matsuda, 2005) and a dataset with shuffled pairs of velocity and activity. Some of the negative control MTA activity levels exhibited some variation from zero, which may be due to measurement bias of the FRET probe. However, the MTA activities were constant near zero for most cases (purple line in **Fig. S4**). We created the shuffled dataset by randomly combining velocity and activity time series in time and location. The MTA activity given by the shuffled dataset was almost zero for all reference velocities and all times (red line in **Fig. S4**). These results indicate that the MTA activity patterns are velocity pattern–specific. Finally, we confirmed that the MTA activity time series (**Fig. 2**) were generalized features in which specific adjacent edges did not predominantly determine the MTA activity (**Fig. S5**). For this purpose, we randomly shifted the time series of adjacent edges to prevent nearby edges from having similar activity time series (**Fig. S5a**). We calculated the MTA activities from the time series of random phases and compared them with the original heatmap. The results revealed no significant difference in MTA activity (**Fig. S5b**), meaning that there was no cluster of adjacent edges that dominate the MTA activity. Therefore, the MTA activities in **Fig. 2** represent the general characteristics of each cell.

### MTA activities are consistent with known individual features

The time series of MTA activity in **Fig. 2** clearly show the specific properties of the respective velocity patterns. Because we displayed all the MTA activity time series shifted to zero at the initial time of the velocity pattern (t = 1), we could evaluate the relative increase and decrease in the activities of the three molecules. First, the MTA activities of all three molecules increased in the “Expansion” pattern (**Fig. 2a**). The rate of increase in activity of Cdc42 and Rac1 was larger than that of RhoA, and RhoA had the largest difference in activity from the other two molecules near the maximum edge velocity. The difference increased with the change in velocity intensity from low to high. In the “Retraction” pattern (**Fig. 2b**), the activities of all molecules decreased from the initial value. The decreasing rates of Cdc42 and Rac1 activities were more significant than that of RhoA. The difference in activities was most significant near the maximum negative velocity and increased with velocity intensity. In addition, during both expansion and retraction, the molecular activities reached their peaks later than the velocity peaks. Therefore, when we calculated the temporal cross-correlation between the velocity and the activity, the coefficient was largest when we shifted the activity time series to the past (see **Fig. 1b**). The result of this time shift to the past is consistent with the results of previous studies that performed temporal cross-correlation analysis (Kunida et al., 2012; Machacek et al., 2009; Tsukada et al., 2008).

The “Acceleration” pattern (**Fig. 2c**), in which the edge accelerates and changes from retraction to expansion for middle and high intensities, showed an activity series similar to that of **Fig. 2a**; the difference between RhoA and the other two increased as velocity intensity changed from low to high. Similarly, in the “Deceleration” pattern (**Fig. 2d**), as shown in **Fig. 2b**, RhoA was more active than the other two, and the difference increased with velocity intensity. Both results in **Fig. 2c** and **d** show that Rho GTPase activities changed smoothly even when the edge reversed its direction of movement, suggesting that at least Rho GTPases do not significantly alter the regulation of expansion and retraction. In contrast to the results in **Fig. 2a** and **b**, the differences between RhoA and the other two activities in **Fig. 2c** and **d** continued to increase. The velocities of “Expansion” and “Retraction” patterns approached zero in the second half, whereas those of the “Acceleration” and “Deceleration” patterns continued to increase. Therefore, it is reasonable that the difference in activities between the Rho GTPases in **Fig. 2c** and **d** continue to increase. Together, the results in **Fig. 2a-d** reveal a balance between the expansion-related molecules Cdc42 and Rac1 and the retraction-related molecule RhoA; this idea is consistent with previous studies (Etienne-Manneville and Hall, 2002; Hall, 1998; Jaffe and Hall, 2005; Pertz, 2010).

The MTA activity time series for constant reference velocities (**Fig. 2e**) provides novel insights for which there are no previous studies to compare. When the edge was at rest (zero velocity), all GTPase activities were at average levels (the activities were standardized to a mean of 0); the activities increased or decreased monotonically for the non-zero constant velocity, whereas the difference between the activities also continued to increase. These results suggest that (1) cells consume energy equivalent to the average GTPase activities in order to hold the edge; (2) there are no velocity-specific GTPase activities or differences between them; and (3) when the activity changes with a constant rate, the distance the edge moves also changes with a constant rate. In order to summarize these results, we need to analyze them using a quantitative mathematical model.

### Formulation of quantitative mathematical models and model selection

To obtain numerical evidence that the three Rho GTPases coordinate to regulate the edge velocity, we formulated a mathematical model that converts MTA activities to edge velocity by considering the results in **Fig. 2** as an aspect of a nonequilibrium open system (Prigogine and Nicolis, 1971). A cell expands and retracts its edges by converting the chemical potential of the active Rho GTPase to low-quality energy. Because cell membrane tension constantly generates internal pressure (Barnhart et al., 2017; Ermilov et al., 2007; Evans et al., 2005; Lieber et al., 2013, 2015), a cell has to produce a mechanical force against internal pressure by consuming energy equivalent to average GTPase activity in order to keep the edge at rest (**Fig. 2e**). When the edge is expanding at a constant velocity, the membrane increases its elasticity according to Hooke’s law, and therefore GTPase activity must further increase in order to overcome the retractive force. Similarly, in constant-velocity retraction, it is necessary to decrease GTPase activity further, because the membrane becomes less elastic. If we approximate the expansion process as an example, both the elastic force and the accumulated chemical potential are proportional to the square of the expansion duration (**Fig. 3a**). This result suggests that tension and GTPase activity mutually change the mechanical equilibrium point. Therefore, we assume that the actin filament, a polymer of ATP-bound actin, is the entity that accumulates the chemical potential of GTPase activity at each time. Rho GTPase activity leads to actin filament formation, and the actin filaments act as scaffolds: the filaments utilize the mechanical force gained from adhesion to the substrate and push back the cell membrane (Footer et al., 2007). Therefore, when the edge is at rest, the tension is in balance with the energy (ATP)-rich actin filaments (**Fig. 3b**; **Fig. S6a**). A cell consumes an average level of active GTPase potential in order to maintain actin filaments. As GTPase activity increases, more actin filaments acquire expansion force from the substrate and expand the edge (**Fig. 3c**; **Fig. S6b** and **c**). In the extending part of migrating cells, filopodia and F-actin are abundant (Genuth et al., 2018; Meyen et al., 2015; Pollard and Borisy, 2003). If the activity returns to the average level, membrane tension will cause the edge to retract (**Fig. S6d**). A reduction in actin filament level leads to weak adhesion to the substrate and causes fast retrograde flow pushed by the membrane (**Fig. 3d**; **Fig. S6e** and **f**). Fast retrograde flow leads to edge retraction (Abe et al., 2018; Burnette et al., 2011; Giannone et al., 2009), decreasing membrane expansion until the membrane returns to mechanical equilibrium. Then, if the activity returns to the average level, the actin filaments will expand their edges (**Fig. S6g**). These cytoskeleton–membrane mechanics have the potential to quantitatively explain the characteristics of the time series of MTA activities at a constant velocity of edge movement (**Fig. 2e**).

**Figure 3.**
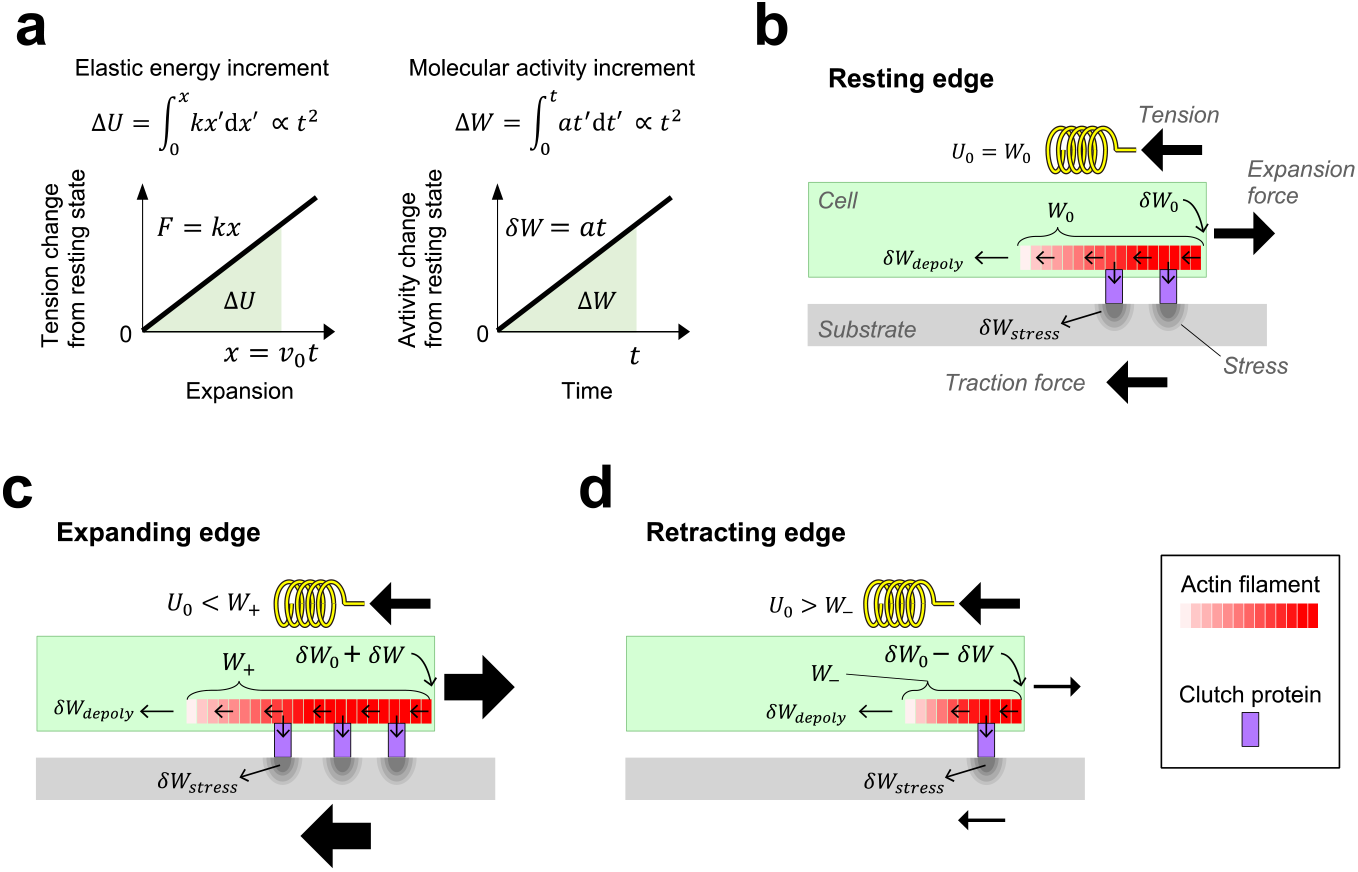
Mathematical modeling based on the chemical potential of GTPases and the elastic energy of membrane tension. **a**. Approximate calculation of energy accumulation for the positive constant velocity case in **Fig. 2e**. As the membrane expands at a constant velocity (*v*_0_), the tension increases at a constant rate, according to Hooke’s law. The elastic energy increment *ΔU* is proportional to *t*^2^ for the expansion duration *t* (left panel). Because the GTPase activity increases at a constant rate (**Fig. 2e**), the increment of stored energy *ΔW* is also proportional to *t*^2^ (right panel). **b**. Energy flow at the resting state of the edge. The chemical energy supply of the average GTPase activity just below the membrane *δW*_0_ forms actin filaments and stores energy (ATP) in them (*W*_0_). Actin filaments play a scaffolding role in generation of traction forces to the substrate and extension forces to the membrane. *U*_0_ is in a state of balance with the elastic energy *W*_0_ of the membrane tension. The actin filaments consume the substrate deformation energy *δW*_*stress*_ through the clutch protein and the ATP hydrolysis energy *δW*_*depoly*_ at the pointed end. Because the sum of these energy consumptions is equal to the supply *δW*_0_, the accumulated energy *W*_0_ does not change. Therefore, if the GTPase activity does not change (*δW*_0_ is constant), *U*_0_ = *W*_0_ is maintained, and the edge continues to be at rest. **c**. Energy imbalance due to increased activity (+*δW*). The edge expands as actin filaments grow and the traction force increases (*W*_+_ = *W*_0_ + *δW* > *U*_0_). As the edge expands, the elastic energy increases, bringing the edge to rest again if *W*_+_ is constant (**Fig. S6**). **d**. Energy imbalance due to decreased activity (−*δW*). Actin filaments decrease, leading to less traction (*W*_−_ = *W*_0_ − *δW* < *U*_0_), and the edges retract. If *W*_−_ is constant, the edge comes to rest again (**Fig. S6**).

Based on the above considerations, a cell requires a change in GTPase activities in order move the edge by taking it out of cytoskeleton–membrane mechanical equilibrium. Therefore, we can write the edge velocity *V*(*t*) at each time *t* as a function *V*(*t*) = *F*(*δW*(*t*)), where *δW*(*t*) is the short-interval change in energy *W* given by GTPase activities. Applying the Maclaurin expansion to this and ignoring terms above the second order, we get

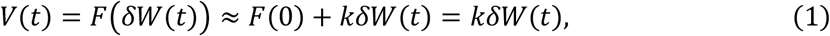

where we set *F*(0) = 0 due to the results in **Fig. 2e** and *k* is a constant parameter. The energy *W* consists of the chemical potentials of the GTPases at each time and the chemical potentials stored in the already formed actin filaments. If *C, R*, and *H* represent the activities of Cdc42, Rac1, and RhoA, respectively, and these variables approximate the change in energy *W* (**Supplementary Note**), we can describe the edge velocity as a linear sum of the GTPase activities and their derivative values as follows:

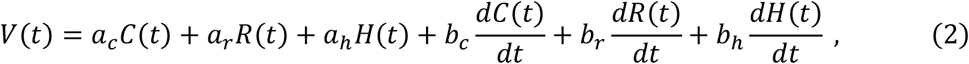

where the six coefficients (*a*_*i*_ and *b*_*i*_; *i* = *c, r, h*) are constants that express the contributions to velocity. Because we have written Eq. (2) in general terms and have not specifically introduced the role of the three GTPase in regulating velocity, this equation represents 2^6^ − 1 (= 63) possible models.

We tested all possible mathematical models using the time series data obtained from the MTA and selected the optimal model by performing model selection and parameter estimation (**Fig. 4a**). Following the general framework of regression, we standardized the variables (mean zero, s.d. one) and minimized the error between the MTA velocity and the model (**Supplementary Note**). We calculated the derivatives by fitting the activity time series with polynomial equations (**Fig. S7**). For model selection, we used the evaluation criteria of the Akaike Information Criterion (AIC) (Akaike, 1973), the Bayesian Information Criterion (BIC) (Schwarz, 1978), and the prediction error of test data by cross-validation (CVMSE). AIC and BIC regress the model on all data; a model with lower error and fewer parameters is better (**Fig. S8a**). CVMSE, which divides the data into training and test data, estimates the parameters on the training data and then selects the model with the lowest prediction error on the test data as the best model (**Fig. S8b**). AIC and BIC avoid models that are overfitted to the data, whereas CVMSE evaluates the prediction performance for unknown data. Models optimized by these criteria represent phenomenological laws that quantitatively decode edge velocities from molecular activities and allow us to quantitatively evaluate the biological role of each Rho GTPase on velocity.

**Figure 4.**
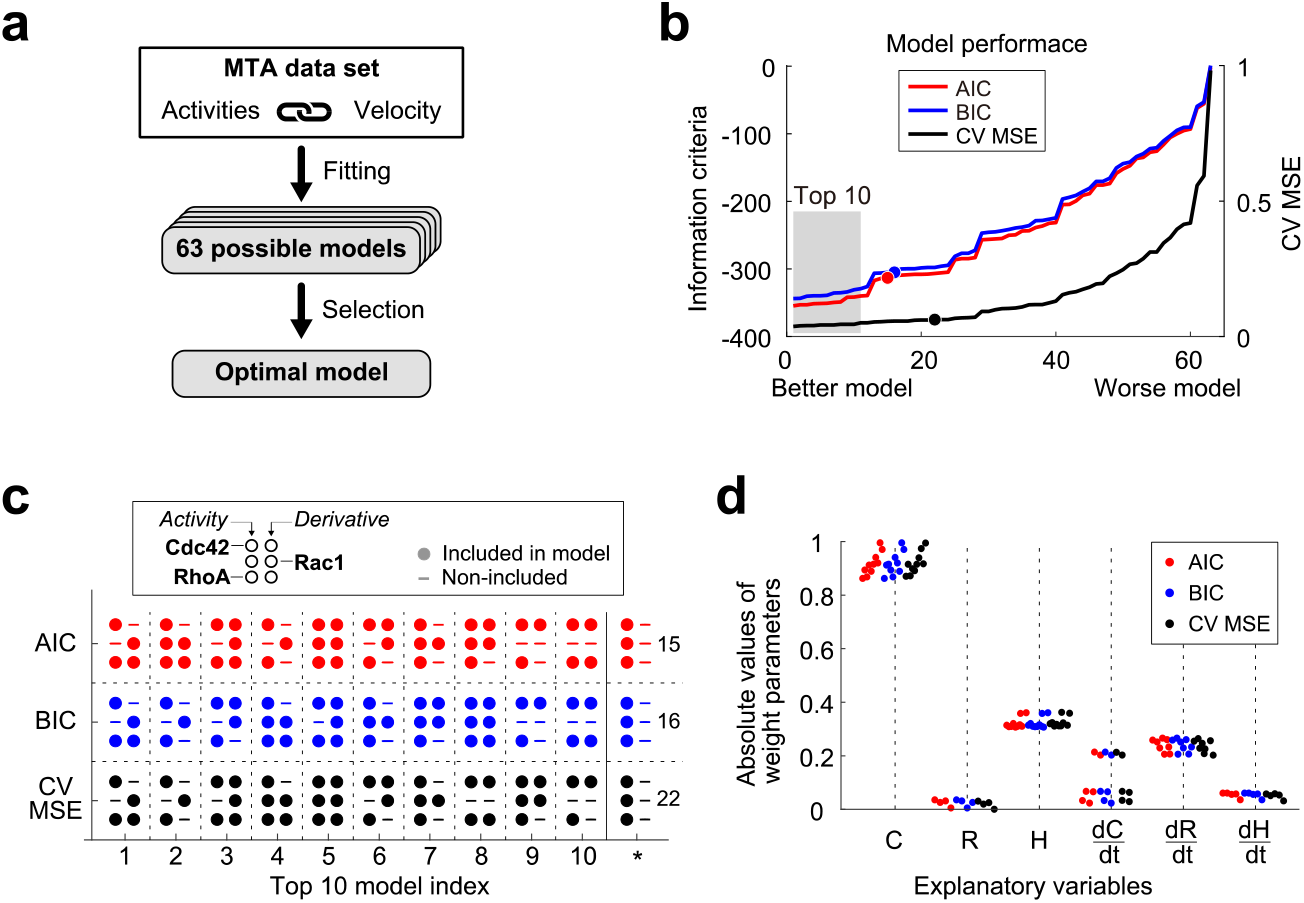
Model comparison using information criterion or cross-validation prediction error. **a**. Schematic procedure for estimating the function for decoding the velocity from the GTPase activities (see **Fig. S8** for details). Applying MTA with five reference velocity patterns, we can obtain pseudo-simultaneous measurements of edge velocity and the activities of three GTPases (Cdc42, Rac1, and RhoA) for each reference. Because the system is unknown, we introduced regression analysis and model selection techniques to estimate the quantitative and phenomenological decoding function from 63 possible models. **b**. Akaike information criterion (AIC, red), Bayesian information criterion (BIC, blue), and prediction error in cross-validation (CVMSE, black) for models with all six variable combinations. We displayed the models sorted in each evaluation; the gray boxes indicate the top ten models. Red, blue, and black circles on the line indicate models with only GTPase concentrations (*C, R*, and *H*). **c**. Comparison of the variables used in the top ten models, where the six variables are represented as 3 × 2 matrices, and the filled circles indicate that they are used in the models (see the inset). The inset on the right column represents the rank of the concentration-only models (circles in **c**). The models that use all variables are the 5^th^, 8^th^, and 5^th^ for AIC, BIC, and CVMSE, respectively. **d**. Comparison of the contribution of each variable in the top ten models (absolute values of the estimated weights). The parameters in the linear regression with standardized variables refer to the weight of the variables. The Cdc42 derivatives split the weights into two groups; the group with the larger weights ranks sixth or lower.

### The optimal model involves three GTPases as variables and predicts unknown data with high accuracy

Because the regression model expressed in Eq. (2) represents all possible roles of the activity variables on velocity, we investigated the most suitable combination for the MTA dataset. We calculated three criterion values (AIC, BIC, and CVMSE) for all 63 models (**Fig. 4b**). The smaller these criterion values are, the better the model is. When we sorted the models in descending order of the value for each criterion, the models were not equally ranked, and the model that regressed only on the GTPase activities without their derivatives did not rank in the top ten (three filled circles in **Fig. 4b**). The top ten models were similar to each other because all included the derivative of the activity (**Fig. 4c**). In particular, many of the models included the activity of Cdc42 and RhoA (*C* and *H*) and the derivatives of the activity of Rac1 and RhoA (*dR*/*dt* and *dH*/*dt*), and a model consisting of these four variables was the best according to all criteria. The contributions of each variable in all top ten models were similar to each other (**Fig. 4d**), and the contributions of the activity of Rac1 (*R*) and the derivative of Cdc42 activity (*dC*/*dt*) were small. Some of the derivatives of Cdc42 had a contribution of ∼0.2, but they were all low-rank models. By contrast, the derivative of RhoA activity (*dH*/*dt*) consistently had a contribution of ∼0.2 to all of the top ten models. Although the top ten models consist of different combinations of variables, these results show that the four variables of the optimal model (*C, H, dR*/*dt*, and *dH*/*dt*) are the keys to decoding the edge velocity.

We next compared the training and testing time series in cross-validation, using the optimal model in **Fig. 4c** (**Fig. 5a**), to determine the performance of velocity decoding. Here, the variables and parameters of the model are displayed, scaled back to the MTA data (see **Supplementary Note**). We performed five-fold cross-validation with five combinations of reference velocities. Hence, the twelve series are the training data (white-background panels in **Fig. 5a**), and the three series are the test data (yellow-background panels in **Fig. 5a**). We found that the model could decode the test velocity time series with significant accuracy in each of five validations. All the parameters obtained for each of the five cross-validations were estimated to be close to each other (**Fig. 5b**). Only the parameter for the activity of RhoA was negative, whereas the others were positive. This observation is consistent with biological findings that RhoA decreases velocity. We could estimate the effect of the derivative of the activity by simply regressing the velocity on the activity series alone (**Fig. S9**). In particular, the errors became more significant for the “Expansion” and “Retraction” reference velocities (**Fig. S9a**), and the Rac1 activity parameter was still almost zero, as shown in the optimal model (**Fig. S9b**). Thus, Rac1 activity does not contribute to slow expansion, but its derivative does contribute to fast edge movement. These results provide numerical evidence for the coordinated regulation of edge velocity by the three Rho GTPases and the novel finding of the contribution of the derivatives of the activities.

**Figure 5.**
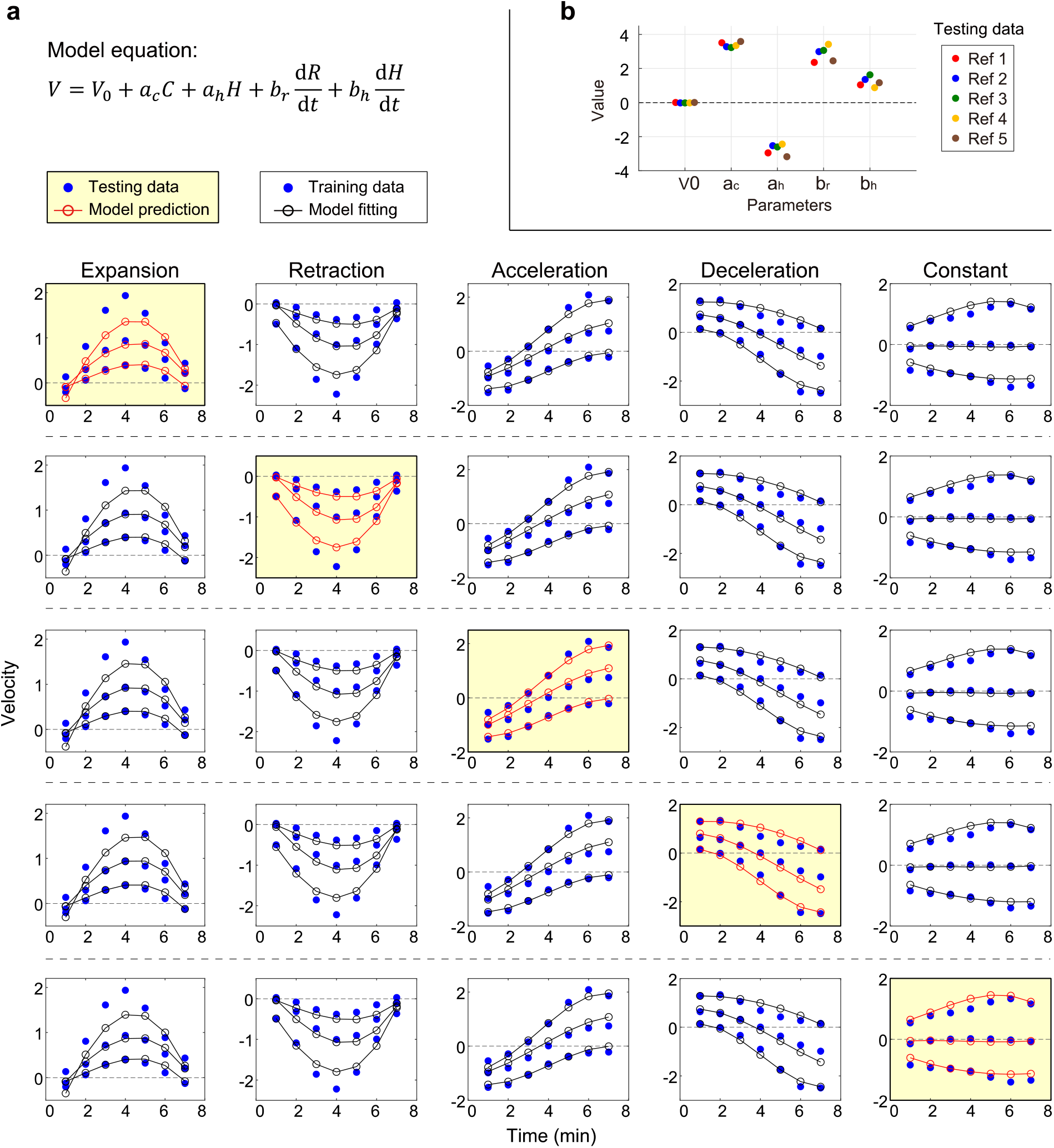
Time series and parameter comparisons in five-fold cross-validation. **a**. Comparison of velocity prediction by the optimal regression model with observed velocity in cross-validation. The regression equation at the top is the optimal model for all AIC, BIC, and CVMSE evaluations. The equation is a formula that returns the standardized variables to the original scale of the variables (see **Supplementary Note**). The blue circle in the graph shows the velocity time series extracted by MTA, the white background shows the training data, and the yellow background shows the test data. We estimated the model parameters from four training data sets (12 series in total), arranged horizontally. The black circles show the time series of the model fitted to the training data, whereas the red circles show the prediction from the molecular activity of the test data. **b**. Comparison of five sets of parameters obtained by cross-validation. Negative numbers mean that an increase in molecular activity causes a decrease in velocity.

## Discussion

### System identification method by combining MTA and regression

Our study was motivated by fundamental difficulties of simultaneous observation of multiple molecules by biosensors and the lack of adequate analytical techniques for individual observation data. Here, we proposed the Motion-Triggered Average algorithm, which averages the activity patterns that trigger the same motion of an edge from the independently observed activities of three GTPases (Cdc42, Rac1, and RhoA). Using a short reference velocity pattern, we could integrate unique and complex movements across multiple cells. Because the cells have similar MTA activity patterns (**Fig. S3**), the cell averages gave reliable activity patterns common to all cells; this allowed decoding of the velocity-time series without overlearning for a particular cell, providing numerical evidence for the three GTPases’ coordinated edge regulation. To check the decodability, it is helpful to regress the velocity on the activities. We have succeeded in formulating the velocity quantitatively using regression with MTA activities. This study is the first to describe coordinated edge velocity regulation based on the activity of three GTPases and decode mechanical phenomena from biochemical signals. In contrast to this study, previous efforts focused on conversion of signals between biochemical molecules (Iglesias and Devreotes, 2012; Janetopoulos et al., 2004; Purvis and Lahav, 2013).

Screening of causes based on specific effects can effectively integrate individual observation and decipher causal inferences of black box targets. In case–control studies, such as genome-wide association studies, individually diagnosed samples are screened for healthy or diseased conditions in order to estimate exposure factors and genes responsible for the disease. Neuroscience requires human subjects to perform identical behavioral tasks in order to measure task-related brain activity. In the machine learning field, supervised learning screens input vectors based on individually assigned labels and reveals the input vectors’ properties. Our proposed MTA is unique and novel in two important ways: extraction of molecular activity time series conditioned on the phenotypic time series of each cell, and extraction of the components common to multiple cells. Because this approach makes it possible to pseudo-simultaneously observe any molecular activities based on phenotypic time series, we can perform phenotype regression with multiple factors. By contrast, regression of edge velocity by a single observed activity time series, rather than by multiple factors or components common to cells, leads to overfitting. In such regressions, the cross-correlation between regression velocity and observed velocity is about the same as that between observed activity and observed velocity (Yamao et al., 2015), and the regression does not improve the information about velocity. When a system of multiple molecules drives cellular function, it is appropriate to use data preprocessing with MTA and regression analysis with the extracted data.

The key to successful analysis with MTA and regression lies in the number of samples in the time series and the validity of the regression equation. To accurately extract the time series of a specific molecular activity by MTA, a sufficient number of samples must be averaged. If there are not enough samples, the accuracy of the MTA will be poor. We sampled a short time series of seven time points to obtain a sufficient number of samples. Although we did not use a large number of cells in this study, this was not an obstacle, because MTA provides high reproducibility. For regression, it is critical to construct a model equation. The optimal model in this study could predict 13 of the 15 velocity time series with reasonable accuracy. However, the errors in the other two series with high acceleration (the maximum intensity in “Expansion” and “Retraction” in **Fig. 5a**) were relatively large, especially around the peak velocity. These large velocities occurred at accelerations or decelerations that exceeded the limits of the velocities that the regression model could represent, and the errors could be due to mechanical factors caused by the deformation of the entire cell that the model did not introduce. Expansion or retraction of the entire cell produces a decrease or increase in local tension (Gauthier et al., 2011). In addition, in this study, we performed a simple least-squares regression without using a method to suppress overlearning such as Ridge regression (Hoerl and Kennard, 1970). However, the small error between the model and the data indicates that there is little noise in the MTA activities and that we could prepare a candidate equation that captures the fundamental structure behind the data. If a large amount of noise persists even after MTA preprocessing, methods such as Ridge regression may be effective. When it is impossible to introduce any biological evidence in the formulation, it may be helpful to use the data to refine the optimal model from a nonlinear polynomial with many terms (Brunton et al., 2016; Schmidt and Lipson, 2009).

### Biological interpretations of the optimal model

Our optimal model indicates that the Cdc42 and RhoA activities are related to the already formed cytoskeleton, and that the derivative of Rac1 and RhoA activities are related to actin polymerization at the barbed end. Based on experimental results, we hypothesize that Cdc42 forms actin bundles to provide a scaffold for the newly formed cytoskeleton to push the membrane, RhoA generates the contraction force between stress fibers, and the derivative activity of Rac1 and RhoA promotes actin polymerization to expand the membrane (**Fig. 6**). Because RhoA has been proposed to perform opposing functions, i.e., expansion and retraction (Martin et al., 2016; O’Connor and Chen, 2013), one possible explanation is that RhoA activity serves to retract and the derivative serves to expand. In addition, the fact that Rac1 does not contribute to slow expansion (**Fig. 5** and **Fig. S9**) is consistent with the idea that the protein is involved in active fast expansion in migration but not in slow expansion such as cell spreading (Steffen et al., 2013). We could interpret this in the same way for the activity derivative of RhoA. It is plausible that the derivatives of the activity time series contribute to the regulation of the edge velocity. Cells are non-equilibrium open systems that undergo state transitions in the process of free energy flow (Prigogine and Nicolis, 1971). Even if the cell does not move, it must consume energy to maintain its morphology and release low-quality energy outside the cell. For other examples, the sodium-potassium pump uses ATP to produce a hyperpolarized state in the membrane potential (Hodgkin and Huxley, 1952); hence, the generation of action potentials is sensitive to the derivative of the current stimulation (Mainen and Sejnowski, 1995) (**Fig. 1c**). In theoretical studies in neuroscience and systems biology, we have shown that signal derivatives may benefit biological functions (Naoki et al., 2005; Sakumura and Ishii, 2006).

**Figure 6.**
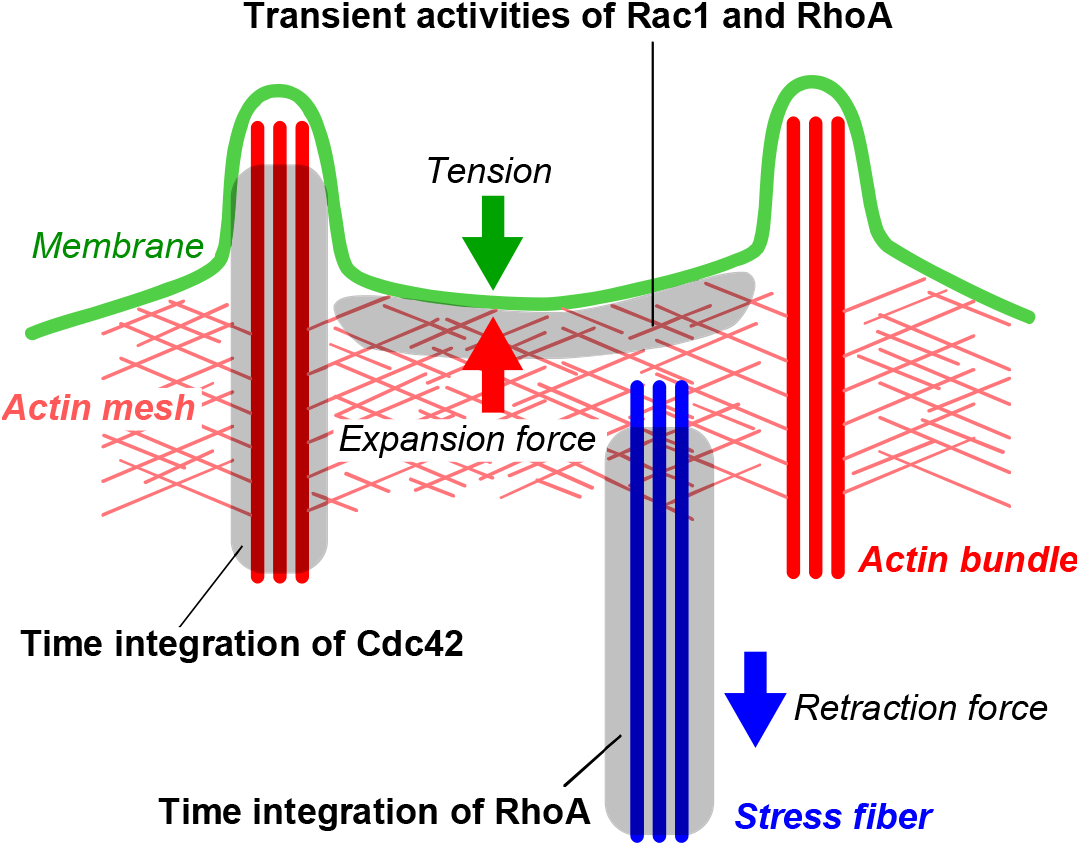
Possible roles of Rho GTPases in the optimal quantitative model. According to the optimal model (**Fig. 3** and **Fig. 5**), the temporal integrals of Cdc42 activity and transient activities of Rac1 and RhoA (red) positively contribute to edge expansion. By contrast, the temporal integrals of RhoA activity (blue) and membrane tension (green) make a negative contribution. Based on our knowledge of the biology, we can interpret Cdc42 integration as actin bundle formation, transient activities of Rac1 and RhoA as actin mesh formation, and RhoA integration as stress fiber formation.

Our optimal model equation provided several clues regarding the mechanism of cell migration. First, the integral of GTPase activity could be related to the decision to migrate. In physical terms, the Cdc42 activity level in the model equation reflects the amount of pre-existing actin filaments (**Supplementary Note**). The amount of actin filament formation reflects the time integration of the actin polymerization signal just below the membrane. This integral would be relevant to decision-making mechanisms in the direction of cell migration (Andrew and Insall, 2007; Baba et al., 2018; Weiner et al., 1999), such as chemotaxis. Even if chemoattractant signals fluctuate, the cell accurately detects the concentration gradient of the signals. One way to achieve stable direction detection is to integrate the concentration of the detected attractants over time to cancel out fluctuations and memorize the gradient information in the actin filaments. This interpretation and the contribution of the derivative of Rac1 activity to fast edge velocity (**Fig. 4** and **Fig. S9**) are consistent with the idea that Cdc42 makes a greater contribution to steering than Rac1 (Yang et al., 2016) and that Cdc42 provides directionality whereas Rac1 provides edge velocity (Beco et al., 2018). Next, our optimal model provided insight into the bistable function (Meinhardt, 1999), which theoretical studies on cell migration often introduce into mathematical models. Because the regression equation for edge velocity is a linear sum of variables related to the molecular activity (**Fig. 5**), the signals represented by Rho GTPases do not effectively undergo nonlinear pathways, such as higher-order reactions, to regulate edge velocity. Even at a constant velocity, the activity does not take on a constant value, such as one of the bistable values (**Fig. 2e**). These results mean that, at least in the cells analyzed in this study, Rho GTPases and their downstream signals do not have the bistable-like features typically seen in nonlinear systems. In that case, can the upstream signals that regulate Rho GTPases have a bistable function? If a bistable system existed upstream, it would monotonically activate or inactivate Rho GTPases, producing time series as shown in **Fig. 2e**. However, because there is an upper and lower limit to the activity level, Rho GTPases cannot continue to increase or decrease their activity, and the expansion and retraction of the edges will eventually terminate. Therefore, even if the factors that act upstream of Rho GTPases have a bistable function, it would not necessarily affect edge expansion or retraction.

In this study, we sought to demonstrate the effectiveness of MTA. As an example, we used time-lapse images of random migration of HT1080 cells. Comparison with migratory cells of other species is not the subject of this study, but MTA analysis using them may yield different activity time series. According to our model based on the mechanical equilibrium between the cytoskeleton and the membrane, the spatial distribution of membrane tension may contribute to directional migration, whereas the membrane tension in the migration direction does not increase. In addition, we can apply the idea of MTA to extract the average time series of any physical quantities that trigger the time series of any phenotype by reusing a huge amount of data that has been obtained previously. We expect that MTA will help to analyze time series of black-box systems.

## Acknowledgments

We thank M. Matsuda, K. Aoki, Y. Inoue, S. Teraguchi, Y. Kumagai, T. Tanaka, and T. Yamada for helpful discussion and technical comments. This work was partially supported by the Japan Society for the Promotion of Science (JSPS) KAKENHI Grant Number (19K20400, K. K.; 20H04283, Y. S.).

Y. S. designed the project; K. K. and N. T. performed the data preprocessing; K. K., Y. S., K. I., and T. N. developed the computational model; and K. K., Y. S., K. I., and T. N. prepared the manuscript.

