## supplemental information for "Decoding cellular deformation from pseudo-simultaneously observed Rho GTPase activities"

**This PDF file includes:**

| | Title |
|---|---|
| Supplementary Figure S1 | Flowchart of motion-triggered average analysis |
| Supplementary Figure S2 | Distribution of assignment of observed time series to reference velocity |
| Supplementary Figure S3 | Cell-to-cell variation of MTA activity time series |
| Supplementary Figure S4 | MTA activities using negative control and randomly shuffled pairs of velocity and activity time series |
| Supplementary Figure S5 | Comparison of MTA activities with time series by random phase segmentation |
| Supplementary Figure S6 | Schematic of the relationship between energy and conversion from GTPase activity change to edge velocity change |
| Supplementary Figure S7 | Derivative time series of MTA activities |
| Supplementary Figure S8 | Model selection algorithms |
| Supplementary Figure S9 | Time series and parameter comparison of five-fold cross-validation with a model using only GTPase activities |
| Supplementary Note | |

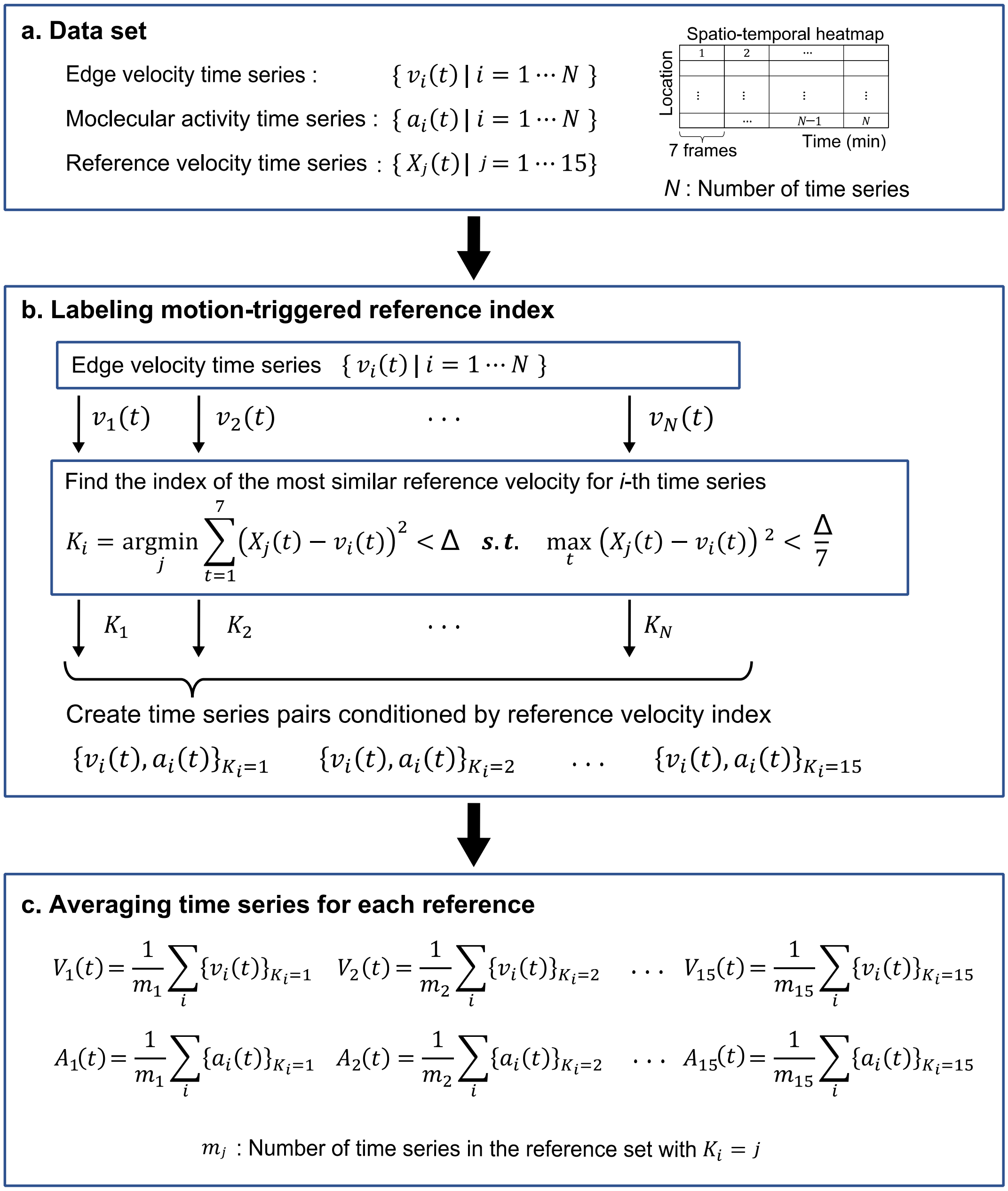


**Figure S1. Flowchart of motion-triggered average analysis.**
**a.** Data preparation step. We created a short time series dataset (seven time frames) from the spatio-temporal heatmaps of edge velocity and GTPase activity (**Fig. 1d**). The indices $(i = 1, \cdots, N)$ are serial numbers of the time series. A total of 15 different reference velocities were prepared, with five patterns ("Expansion," "Retraction," "Acceleration," "Deceleration," and "Constant") with three intensities each $(j = 1, \cdots, 15)$. **b.** Labeling step. We calculated each velocity time series error with the reference velocity time series in order to label the index of the reference velocity with the smallest error. The tolerance parameter $\Delta$ is 10. By classifying the time series for each of those indices, we obtained 15 sets of paired time series of velocity and activity. Most of the time series were assigned one of the 15 reference indices (**Fig. S2**). **c.** Averaging step. We averaged the time series of the velocity and activity for each index of the reference velocity.

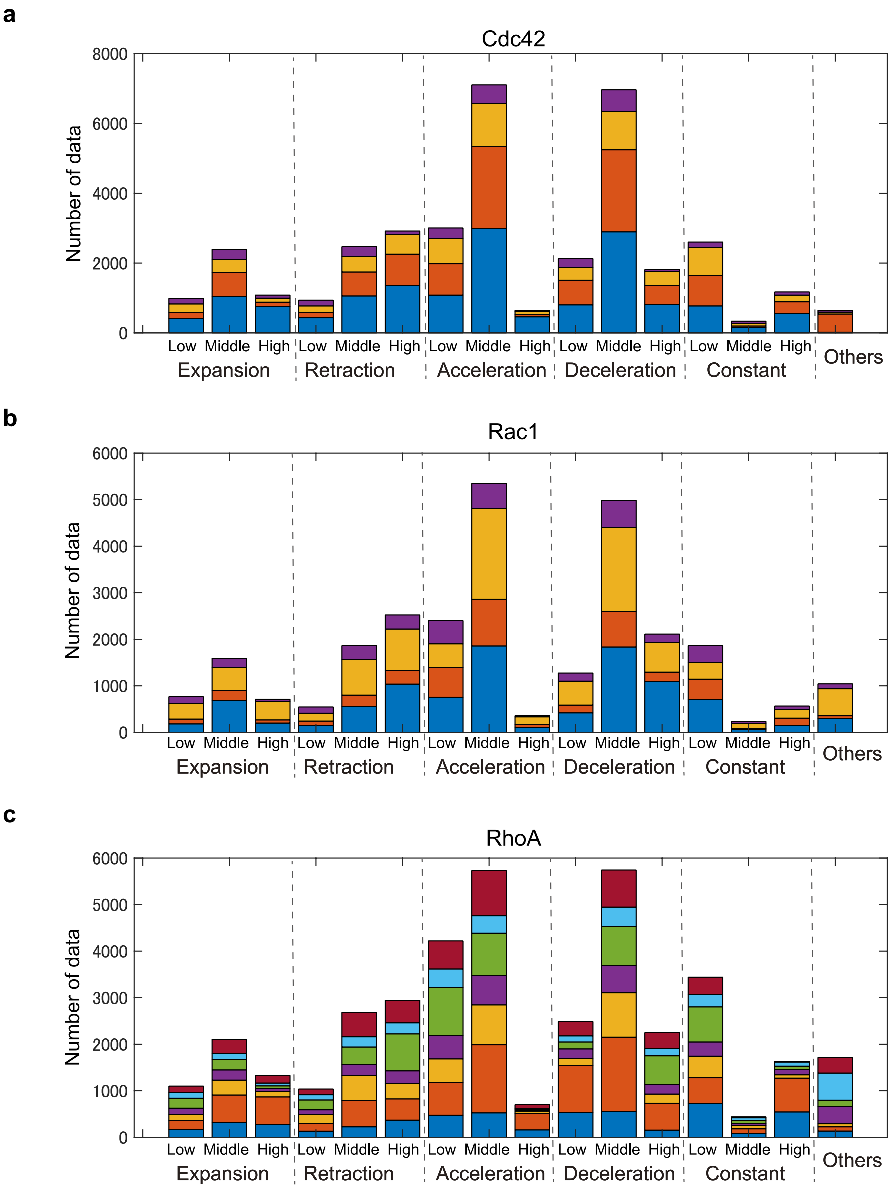


**Figure S2. Distribution of assignment of observed time series to reference velocity.**
For each of the 15 reference velocities and the other patterns, the number of time series assigned by the procedure in **Fig. S1** is shown in different colors for each cell (**a.** Cdc42, n = 4; **b.** Rac1, n = 4; and **c.** RhoA, n = 7).

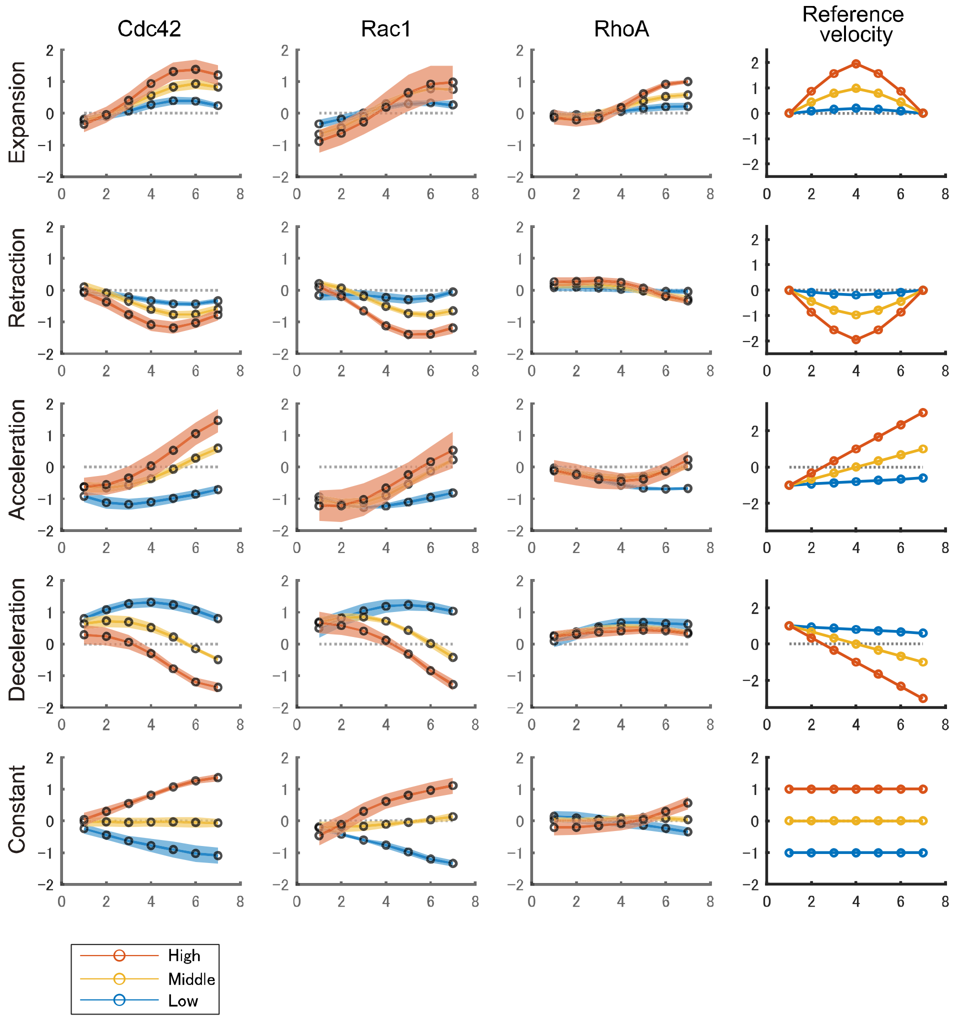


**Figure S3. Cell-to-cell variation of MTA activity time series.**
The color bands represent the standard error of the MTA (black circles), indicating the reliability of the mean. The right column displays the reference velocity patterns for each row in the corresponding color. The number of cells is n = 4, 4, and 7 for Cdc42, Rac1, and RhoA, respectively.

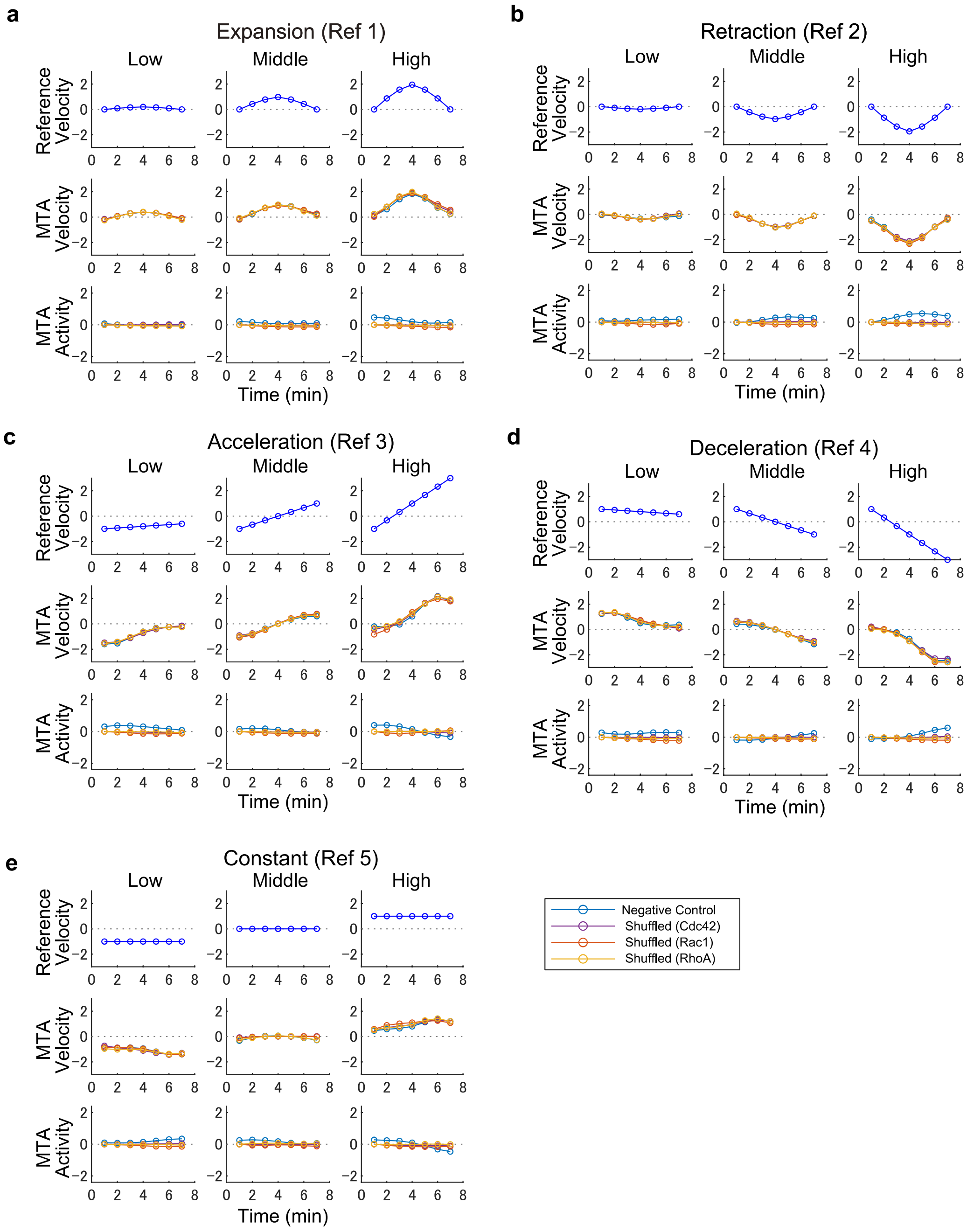


**Figure S4. MTA activities using negative control and randomly shuffled pairs of velocity and activity time series.**

Same as Fig. 2, but cyan lines represent MTA performed with images observed using probes unrelated to the velocity control (Kurokawa and Matsuda, 2005) (n = 7). Because the FRET intensity is in principle uninformative for the 15 reference velocities (**a-e**), the average activity is zero in most cases. Some probes deviate from the mean of zero due to probe artifacts. The other three colored lines represent MTA with randomly shuffled pairs of velocity and activity time series, which are uninformative for the 15 reference velocities.

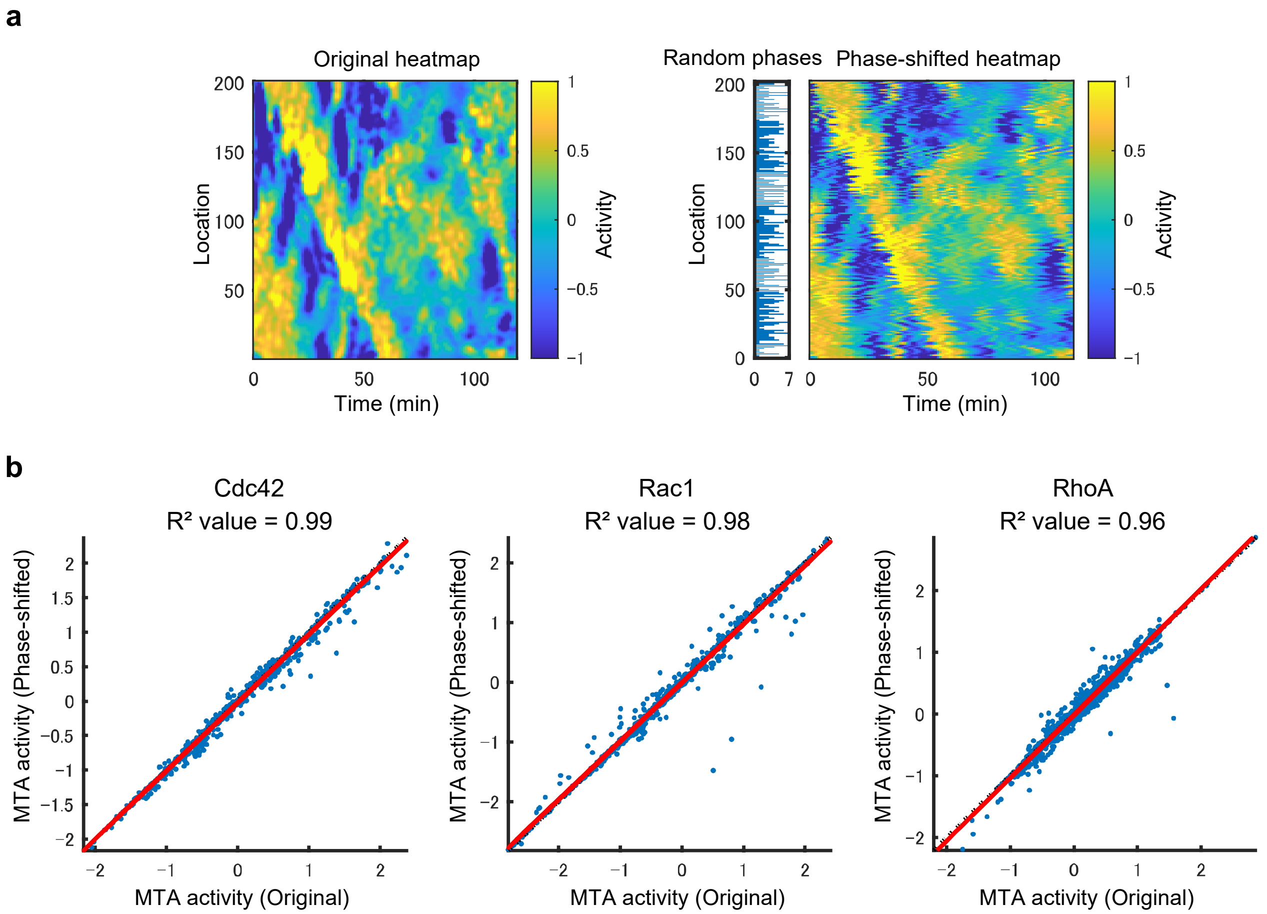


**Figure S5. Comparison of MTA activities with time series by random phase segmentation.** **a.** Sample heatmaps of original and randomly phase-shifted time series. The left figure shows the original heatmap. The figure on the right shows the phase-shifted heatmap with random phases for each location represented by horizontal bars. We sampled phases randomly from 1 to 7 to avoid having adjacent edges with similar time series. **b.** Comparison of MTA activity from the original heatmap with the activity from the phase-shifted heatmap. There are 105 extracted activities from five reference velocity patterns, three velocity intensities, and seven observation time points per cell (n = 4, 4, and 7 for Cdc42, Rac1, and RhoA, respectively).

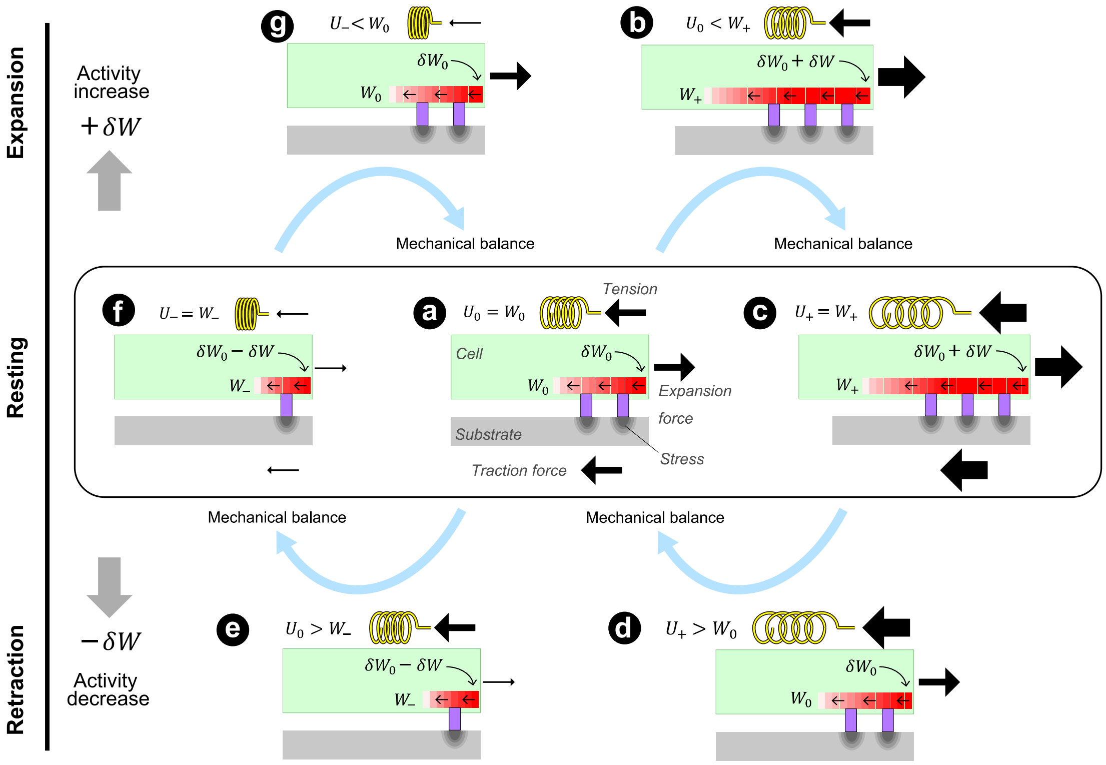


**Figure S6. Schematic of the relationship between energy and conversion from GTPase activity change to edge velocity change.**

The figures in the middle row represent the resting state at various mechanical balances of membrane expansion and tension. The upper and lower rows represent the state transition in the expansion direction due to the activity increase and the state transition in the retraction direction due to the activity decrease, respectively. **a.** The mechanical balance with the average activity, $\delta W_0$. The stored energy $W_0$ balances with the elastic energy $U_0$ (Fig. 3a). **b.** When the activity increases, the actin filament stores more energy to generate an expansion force ($U_0 < W_+$). **c.** The passive membrane increases tension following the expansion force and restores mechanical balance ($U_+ = W_+$). **d.** If the activity returns to the average level ($U_+ > W_0$), the membrane retracts until the mechanical balance disappears (**a**). **e.** When the activity level decreases further and falls below the average level, the expansion force weakens ($U_0 > W_-$), and the membrane tension causes retraction. **f.** As retraction progresses, a weak mechanical balance is established ($U_- = W_-$) and the edge comes to rest. **g.** When the activity level recovers to the average level, the cell gains an expansion force stronger than tension ($U_- < W_0$) and regains the average level of mechanical balance.

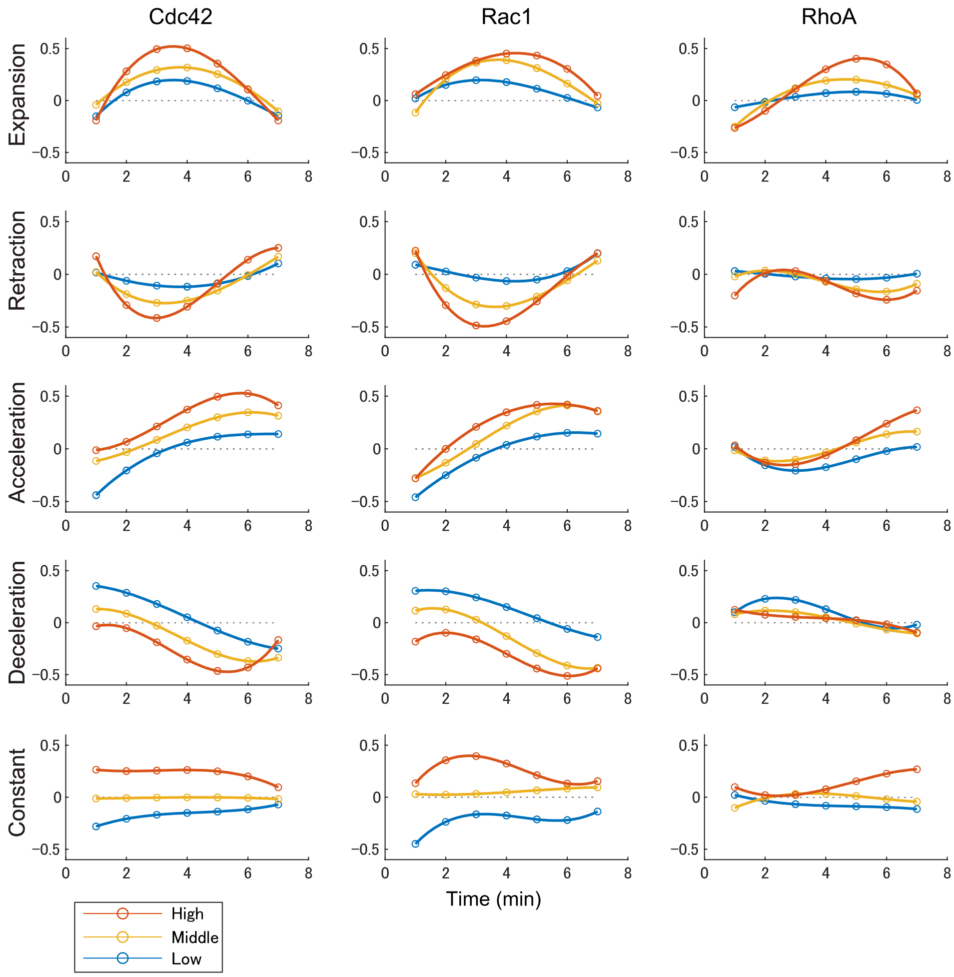


**Figure S7. Derivative time series of MTA activities.**
We obtained the derivatives of the activities by fitting the activity time series in **Fig. 2** with a third-order polynomial.

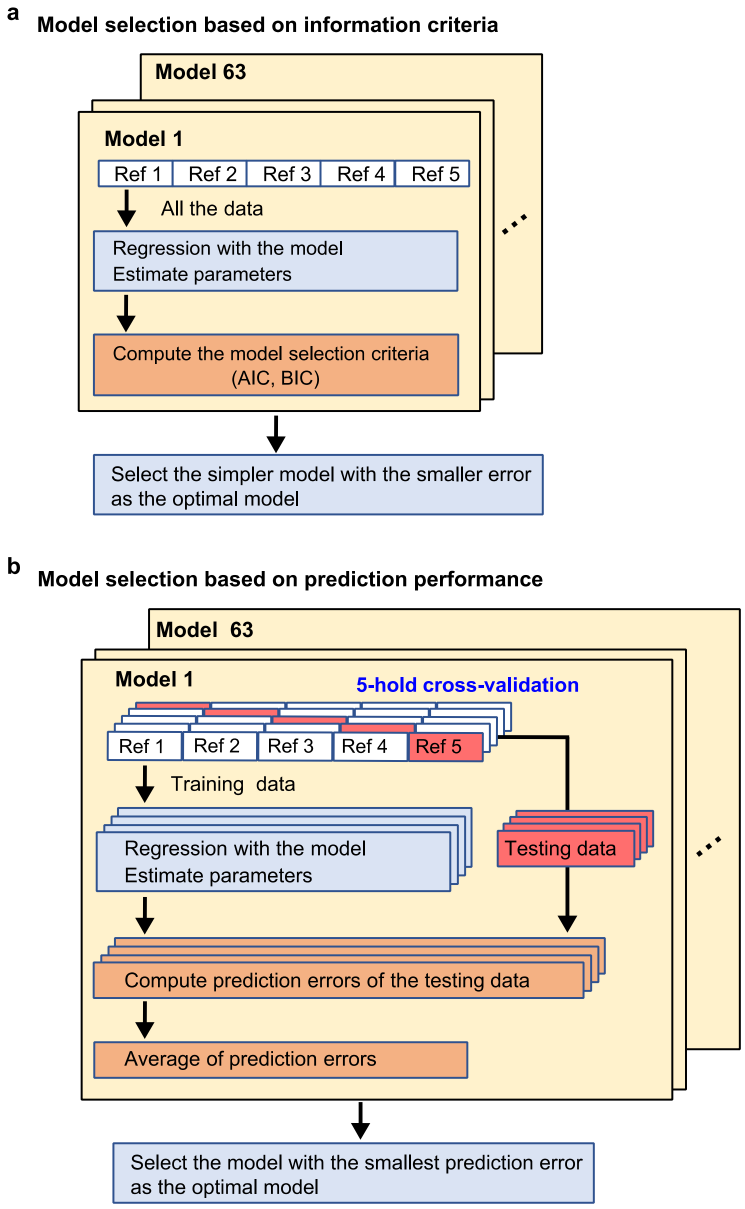


**Figure S8. Model selection algorithms.**
**a.** Schematic of the model selection procedure based on the information content criterion. A generalized model is selected by fitting the model with all the data and satisfying the trade-off condition of small error and few parameters (AIC or BIC). **b.** Schematic of the procedure for model selection based on prediction error against the testing data in cross-validation. The optimal model is best able to predict untrained data.

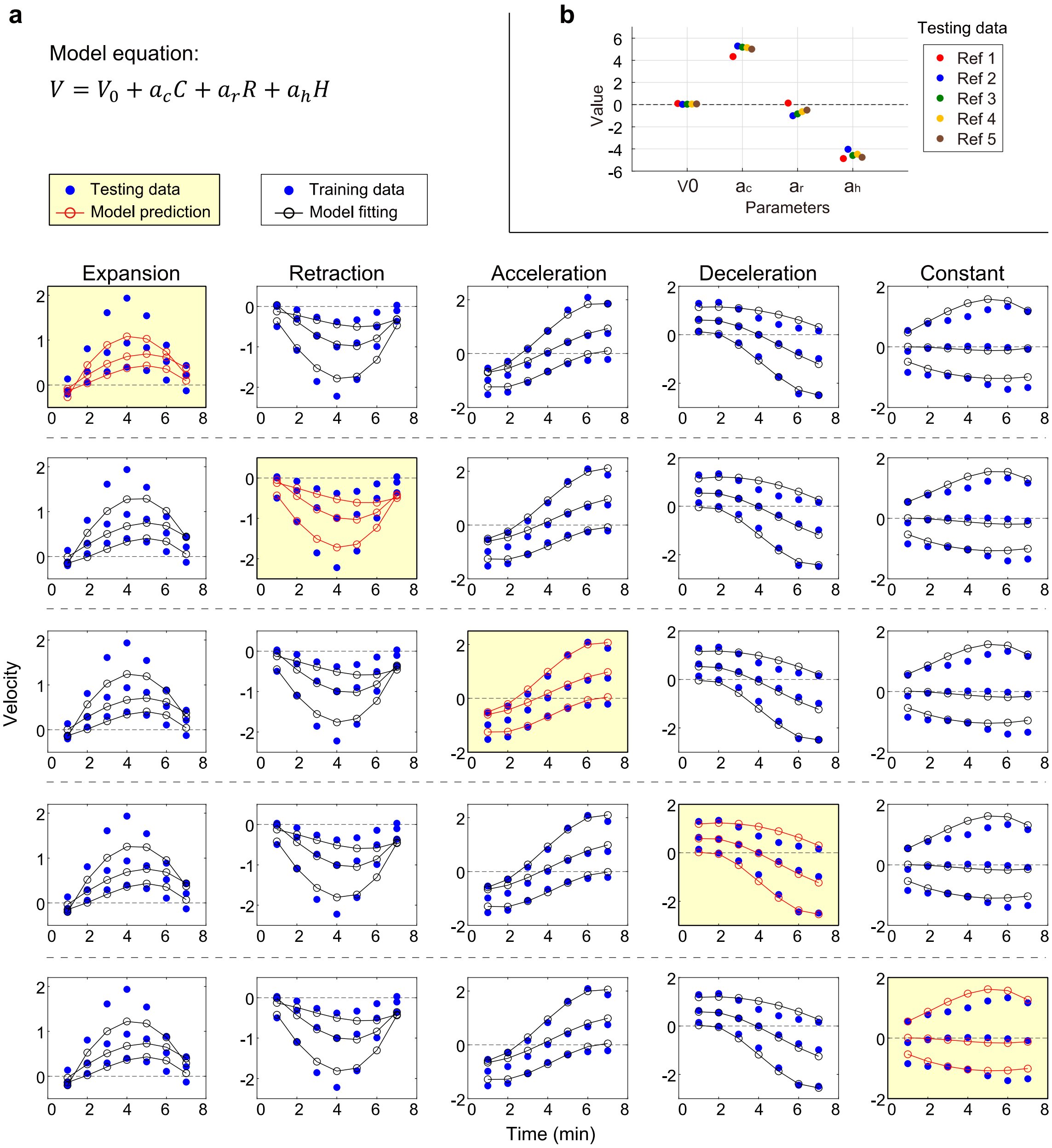


**Figure S9. Time series and parameter comparison of five-fold cross-validation with a model using only GTPase activities.**

**a.** Regression equation composed of GTPase activities without their derivatives. The blue circle in the graph shows the velocity time series extracted by MTA, the white background shows the training data, and the yellow background shows the test data. We estimated the model parameters from four training data (12 series in total) arranged horizontally. The black circles show the time series of the model fitted to the training data, whereas the red circles show the prediction of the molecular activity of the test data. **b.** Comparison of five sets of parameters obtained by cross-validation. Negative numbers mean that an increase in molecular activity causes a decrease in velocity.

### Supplementary Note

#### 1. Time-series data set of edge velocity and Rho GTPase activities

In this study, we used spatiotemporal quantitative data of edge velocity and Rho GTPase activity during spontaneous cell migration obtained in our previous work (Kunida et al., 2012). The activity of Rho GTPase was measured using a FRET biosensor. The plasmids encoding FRET biosensors were as follows: Raichu-Rac1/1011x and pRaichu-Cdc42/1054x (Itoh et al., 2002), pRaichu-RhoA/1294x (Yoshizaki et al., 2003), and pRaichu-Pak-Rho/1110x, which is a negative control (Kurokawa and Matsuda, 2005). We segmented the spatiotemporal quantitative data at seven time points and used the time series data of edge velocity and molecular activity for each locational segment as the analysis data set.

#### 2. Derivation of the regression equation for edge velocity

If we denote the energy given by the activities of Rho GTPases as W, as described in the text, W is a function of the chemical potentials of the GTPases (Cdc42: $C$、Rac1: $R$、RhoA: $H$) and the chemical potentials stored in the actin skeleton. First, we express the stored energy in terms of activity potential. If we approximate the actin skeleton as a one-dimensional bundle structure or a two-dimensional sheet structure with average fixed width, the bundle or sheet will form at a rate proportional to the GTPase activity at each time point. Therefore, we can express the amount of actin filament formed by each GTPase as the time integral of the activity, $S_C(t) = \int_{t_0}^{t} G(t,t')C(t')dt'$, $S_R(t) = \int_{t_0}^{t} G(t,t')R(t')dt'$, and $S_H(t) = \int_{t_0}^{t} G(t,t')H(t')dt'$, where the subscripts represent the type of GTPases. The function $G(t,t')$ is a function that decays depending on the time difference $t - t'$ and represents the effect of depolymerization of actin. In the case of

$t = t'$, $G(t,t)$ is the constant $G_0$. Because the subsequent process of cytoskeletal formation is primarily mechanics-based, the six variables of activity and their integrals can be considered to act on the membrane expansion in parallel and independently.

We can write the energy supply $\delta W$ from the GTPase activities during the small time interval $\delta t$ as

$$\delta W = W(t + \delta t) - W(t) \approx \frac{dW}{dt}\delta t, \tag{S1}$$

by approximation using the Taylor expansion. The total derivative of $W$ with time is the sum of the partial derivatives of each variable:

$$\frac{dW}{dt} = \sum_{y=\{S_c,S_r,S_h\}} \frac{\partial y}{\partial t}\frac{\partial W}{\partial y} + \sum_{x=\{C,R,H\}} \frac{\partial x}{\partial t}\frac{\partial W}{\partial x}. \tag{S2}$$

Substituting this into Eq. (S1), we get

$$\delta W \approx \sum_{y=\{S_c,S_r,S_h\}} \frac{\partial y}{\partial t}\frac{\partial W}{\partial y}\delta t + \sum_{x=\{C,R,H\}} \frac{\partial x}{\partial t}\frac{\partial W}{\partial x}\delta t. \tag{S3}$$

According to the energy conservation law, because the energy $W$ is a linear function of the molecular activity energy ($x$) and the stored energy in the cytoskeleton ($y$), the derivative of $W$ by these variables will be constant. Therefore, Eq. (S3) can be written as,

$$\delta W \approx \sum_{x=\{C,R,H\}} \left(G_0 w_x^{'} \delta t\right) x + \sum_{x=\{C,R,H\}} (w_x \delta t)\frac{dx}{dt}, \tag{S4}$$

where $w_x$ and $w_x'$ are constants, and the partial derivative $\partial y/\partial t$ is replaced with $G_0 x$. In addition, $\partial x/\partial t = dx/dt$ because the GTPase activity time series in this study are timedependent observations. Finally, by replacing the constants, we get

$$\delta W \approx A_c C + A_r R + A_h H + B_c \frac{dC}{dt} + B_r \frac{dR}{dt} + B_h \frac{dH}{dt}. \tag{S5}$$

### 3. Regression procedure

**Data preprocessing:** It is necessary to conduct regressions with the standardized objective and explanatory variables. The data quantified from the experiment is a time series of seven time points of the velocity of the cell edge and the FRET ratio of GTPases around the edge. We estimated the activity derivatives by regressing the series of FRET ratios with a cubic function (**Fig. S7**). In order to regress velocity using different physical units of activity and their derivatives, we standardized each observation (velocity, activity, and derivative of activity) as follows:

$$x = \frac{X - \mathrm{mean}(X)}{\mathrm{std}(X)}, \tag{S6}$$

where $x$ is the standardized data of $X$.

**Regression Equation and Optimization:** Using the standardized data set, we estimate the parameters that satisfy the following regression equation:

$$v = p_c c + p_r r + p_h h + q_c \dot{c} + q_r \dot{r} + q_h \dot{h}\,, \tag{S7}$$

where $c,\ r,\ h,\ \dot{c} = dc/dt\,,\ \dot{r} = dr/dt\,,$ and $\dot{h} = dh/dt$ are standardized variables for each variable and $\theta = \{\,p_i,\ q_i \mid i = c,\ r,\ h\,\}$ are parameters indicating the importance of each variable. Let the experimentally observed edge velocity be $V_{ex}$ and its standardized value be $v_{ex}$. The optimal parameter set $\theta_{opt}$ that minimizes the error between the standardized velocity and the regression velocity was determined as follows:

$$\theta_{opt} = \arg\min_{\theta} \sum_{t \in D} \| v_{ex}(t) - v(t) \|^2 , \tag{S8}$$

where $D$ represents the set of time points of the edge velocity fitting the model.

**Scale transformation of regression equation:** Equation (S6) linearly transforms the observed and standardized data so that we can convert the standardized data to the observed data:

$$V = V_0 + a_c C + a_r R + a_h H + b_c \dot{C} + b_r \dot{R} + b_h \dot{H} , \tag{S9}$$

where $\dot{C} = dC/dt$、$\dot{R} = dR/dt$、$\dot{H} = dH/dt$.
